# Continuous high-resolution *in vivo* imaging reveals tumor-specific dissemination in an embryonic zebrafish xenograft model

**DOI:** 10.1101/215921

**Authors:** N Asokan, S Daetwyler, SN Bernas, C Schmied, S Vogler, K Lambert, M Wobus, M Wermke, G Kempermann, J Huisken, M Brand, M Bornhäuser

**Affiliations:** Center for Regenerative Therapies Dresden (CRTD), Technische Universität Dresden, Germany; Division of Hematology, Oncology and Stem Cell Transplantation, Department of Medicine I, University Hospital Carl Gustav Carus, Technische Universität Dresden, Germany; Max Planck Institute of Molecular Cell Biology and Genetics (MPI-CBG), Dresden, Germany; German Center for Neurodegenerative Diseases (DZNE) Dresden, Germany; National Center for Tumor Diseases (NCT), Dresden, Germany; German Consortium for Translational Cancer Research (DKTK), DKFZ, Heidelberg, Germany

## Abstract

Mechanisms mediating tumor metastasis are crucial for diagnostic and therapeutic targeting. Here, we take advantage of a transparent embryonic zebrafish xenograft model (eZXM) to visualize and track injected human leukemic and breast cancer cells in real time using selective plane illumination microscopy (SPIM) for up to 30 hours. Injected cells exhibited disease-specific patterns of intravascular distribution with leukemic cells moving faster than breast cancer cells. While breast cancer cells predominantly adhered to nearby regions, about 30% invaded the avascularized tissue, reminiscent of their metastatic phenotype. Survival of the injected tumor cells was partly inhibited by the cellular innate immune system of the recipient embryos and leukemic cell dissemination was effectively inhibited by pharmacological ROCK1 blockade. These observations, and the ability to image several embryos simultaneously, support the use of eZXM and SPIM imaging as a functional screening platform to identify compounds that restricts cancer cell spread and invasion.

---

Tumor metastasis is a highly dynamic, complex, and multistage process during which primary tumor cells disseminate from their site of origin, intravasate, and then leave the blood stream to invade distant organs^1,2^. Metastases represent a major determinant of cancer-associated morbidity and death, and an in-depth understanding of the steps involved in this process would lead to improved diagnostic and therapeutic regimens. Even though several in *vitro* models have been developed to dissect the metastasis cascade, they do not adequately mimic the in vivo complexity of this process. Existing in vivo models and various in vivo imaging tools, such as intra-vital imaging^3–8^, have played a pivotal role in studying certain facets of metastatic spread but have inherent limitations with respect to their translational relevance as they are mostly based on murine tumors or artificial xenotransplants. Nonetheless, these studies have provided unique insights into the mechanisms of tumor invasion by capturing migration, plasticity of single cells and their tumor microenvironments, and associated changes in gene expression. Although these advances have enabled a better understanding of metastasis, existing models are not amenable to visualization and continuous monitoring of tumor cells in real time.

The zebrafish is increasingly used as model system to understand tumor progression and dissemination^9,10^, and specific advantages such as optical clarity during embryogenesis, availability of pigment-deficient zebrafish^11^, and amenability to transplantation assays^12^ make the zebrafish a versatile animal model for cancer research^13–15^. Importantly, gene expression profiles of human and zebrafish cancers, such as liver cancer, leukemia (T-ALL), and melanoma, show a high degree of similarity, suggesting evolutionary conservation of pathways associated with cancer progression^16,17^. Also, various human tumor cell lines, including melanoma, glioma, hepatoma, lung cancer, pancreatic cancer, ovarian carcinomas, breast cancer, prostate cancer, retinoblastoma, and leukemia, have been xenotransplanted into zebrafish^18^ to study several aspects of tumorigenesis, like tumor cell migration, angiogenesis, extravasation, and micrometastases^19–23^. Most importantly, proof-of-concept studies have suggested that xenogeneic tumor transplant models using zebrafish embryos can be used as a screening platform to identify novel therapeutic compounds and approaches^24–27^. However, long-term, high resolution, time-lapse images of such transplanted tumor cells are lacking, and their behavior in circulation has not been continuously monitored. This information would be very valuable and help analyze the dynamics of tumor cell spread and invasion in real-time.

Therefore, to understand the dissemination and behavior of tumor cells in circulation, we utilized an existing embryonic zebrafish xenograft model (eZXM). Here, we describe the use of high-resolution, non-invasive, selective plane illumination microscopy (SPIM)^28^ to visualize the characteristics of tumor cells in real time and investigate the metastatic process in *vivo.* The SPIM images revealed adaptations in tumor cell morphology in accordance with their surrounding environment. Using an in-house developed, semi-automated tracking method, we could identify distinct intravascular migration patterns and provide, for the first time, tumor-specific speed and distance measurements. Further, we show that the transplanted tumor cells undergo extravasation and invade the host embryo. Finally, we also demonstrate the suitability of our model for therapeutic intervention using the anti-leukemic drug, Fasudil, a ROCK1 inhibitor. These aspects, combined with the versatility in imaging techniques, make the zebrafish an ideal platform for direct and continuous *in vivo* observation of tumor cells to enable a better understanding of tumorigenesis.

## Results

### Dissemination of tumor cells in vasculature

We utilized early stage embryos (48 hpf) of the *Tg(kdrl:EGFP)*^*s843*^ zebrafish reporter line with green fluorescent vasculature^29^ on a *casper* background, for in *vivo* visualization of engraftment^11^. SPIM^30^ was used to monitor and characterize the dissemination profiles of the triple negative breast tumor cell line MDA-MB231 (representative solid tumor-cell xenograft) and a leukemic cell line OCI-AML3_eGFP (representative leukemic xenograft) in real-time. Freshly isolated human hematopoietic stem and progenitor cells (hHSPCs) from healthy donors were used as controls. Time-lapse images were acquired for up to 30 hours for all these cell types and analyzed. Interestingly, we found that irrespective of the tumor cell type injected, all cells disseminated similarly throughout the embryo from head to tail (DoC; Suppl_movie_1, 2 & 3; Fig_1a). After 5 hpi, we observed that while the solid tumor cells preferred to adhere to nearby regions, leukemic cells tended to migrate continuously. To understand the migration patterns and morphological changes in individual cells, higher magnification time-lapse images of injected embryos were acquired. These images revealed that in vivo, irrespective of cell type, tumor cells migrated as a combination of individual cells, cell streams, or clusters (only breast cancer cells), and that only few cells stayed non-motile and became adherent after homing to one site (Fig_1b). Higher magnification images also revealed that solid tumor cells (breast cancer) showed an amoeboid type of migration (especially in the intersegmental vessels), as they squeezed themselves into vessels (Fig_1c). These cells also formed large protrusions with filopodia-like structures at the trailing end. However, in other parts of the embryos these breast cancer cells were found to be round and more compact in shape. In contrast to solid tumor cells, leukemic cells and hHSPCs were mostly found to be round in shape.

**Fig_1.**
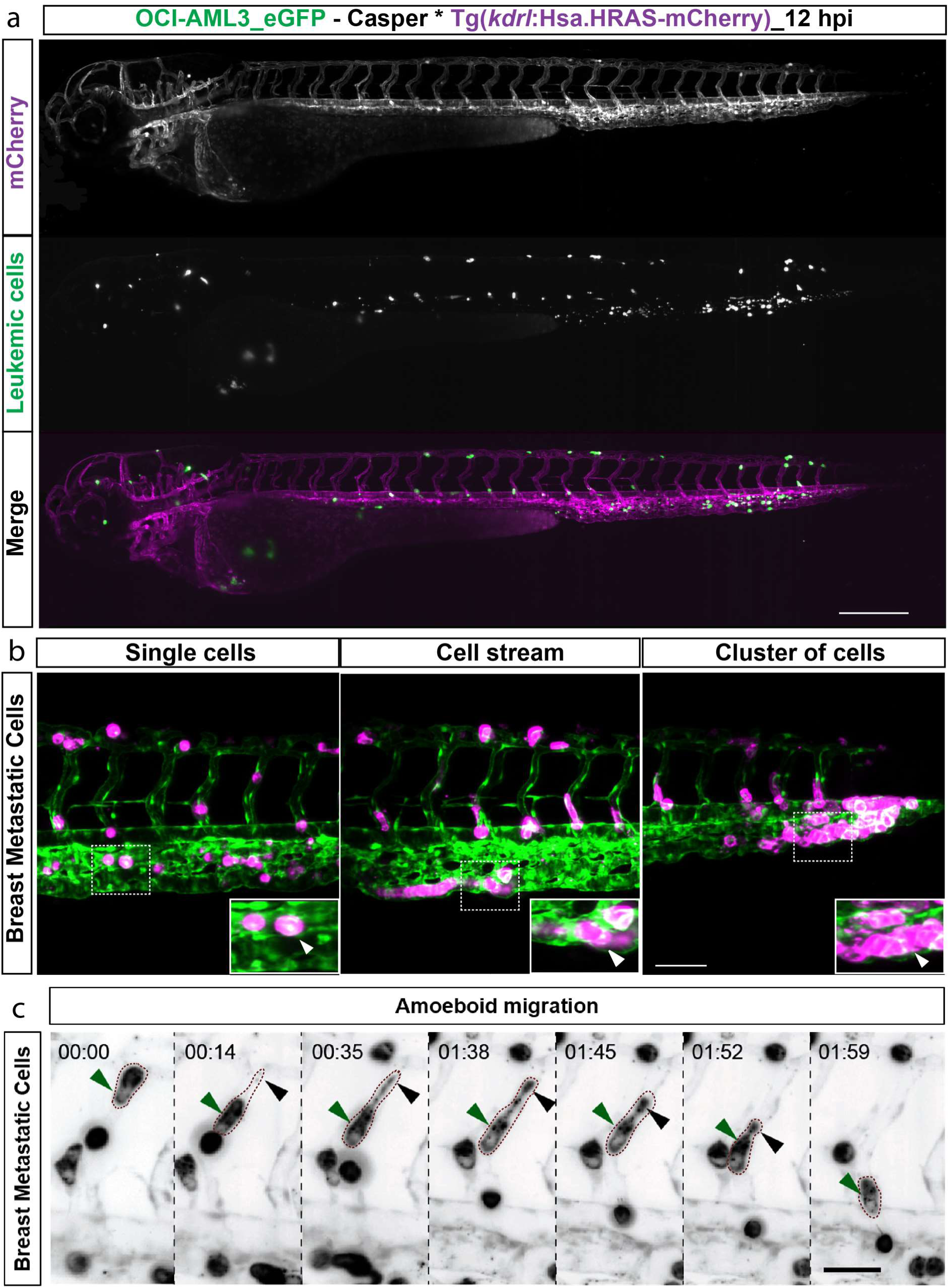
Dissemination and migration modes of tumor cells. (a) Snapshot from time-lapse movie of eZXM expressing the vascular marker *Tg(kdrl:Hsa.HRAS-mCherry)* in *casper* background injected with eGFP labeled leukemic cells (OCI-AML3_eGFP). The cells disseminated throughout the embryo. Vasculature in green, leukemic cells in magenta; scale bar: 500 μm. (b) High-magnification SPIM revealed diverse migratory modes of breast tumor cells. Representative images of tumor cells migrating either as single cells (left), loosely attached cell streams (center), or cluster of cells (right) indicated by white arrowheads. Insets showed the higher magnification of dotted boxes, scale bar 100 μm. (c) A breast tumor cell (MDA-MB-231, green arrowheads) migrating through an intersegmental vessel in an amoeboid fashion (as indicated by dashed brown border). The cell formed a large protrusion, with a filopodia-like arm at the trailing end (black arrowheads). Time shown as h:min, scale bar 50 μm.

### Leukemic cells display rapid intravascular dissemination

Direct in *vivo* recording of SPIM data from the eZXM allowed us to quantitatively analyze parameters of cell dissemination. We applied an in-house developed tracking method for the cells in anterior-posterior and dorsal-ventral direction, combined with manual correction of the tracking results. Dissemination characteristics were described in terms of maximum and total distance travelled, and net distance. Maximum distance travelled was defined as the largest distance between any two given time points in the cell’s migratory path, net distance as the distance separating a cell’s first (origin) and last (final) positions over the entire movie of 30 hours, and the total distance travelled as the path taken by the cells from their origin to their final position (Fig_2a).

**Fig_2.**
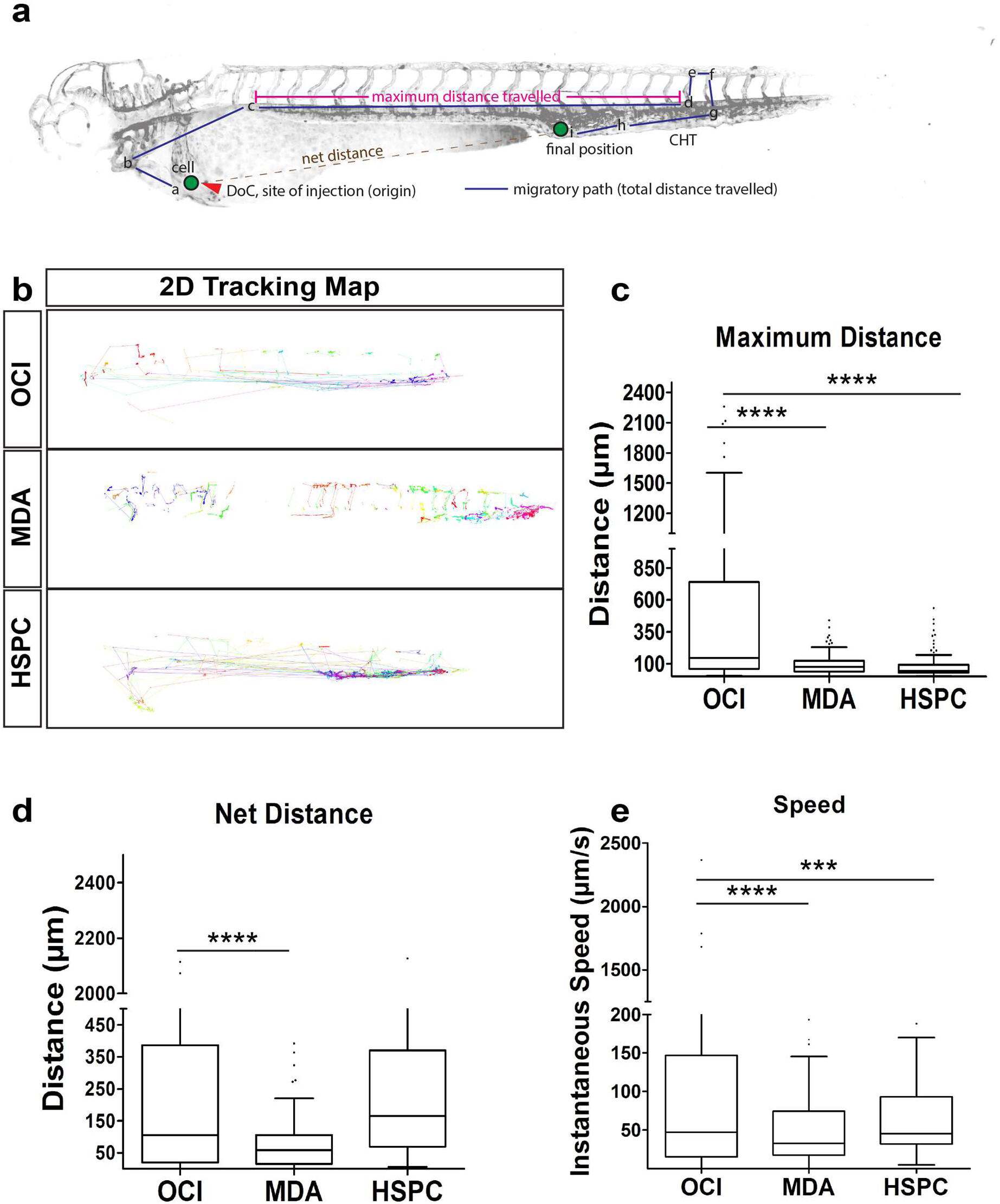
Tumor cell tracking revealed dynamics of xenografted cells. (a) Schematic of the quantified dissemination characteristics of a xenografted cell (green). DoC Duct of Cuvier, CHT caudal hematopoietic tissue. (b) 2D tracking map of the migratory path taken by every cell in *vivo* using semi-automated tracking analysis of the SPIM time-lapse movies. The tracking map revealed circulatory paths of leukemic cells (OCI, top), short migratory paths of breast tumor cells (MDA, center), and mixed migratory patterns of hematopoietic stem and progenitor cells (HSPC, bottom). Each color represented an individual cell. (c-e) Representative plot of R-analysis of the cell tracking revealed that (c) the maximum distance between any two time points was higher in leukemic cells (OCI) compared to either breast tumor cells (MDA) or HSPCs. (d) The net distance was significantly higher for leukemic cells (OCI) compared to breast tumor cells (MDA), but comparable to HSPCs. (e) Intravascular speed measurements in all three cell types: breast tumor cells (MDA), leukemic cells (OCI) and HSPCs revealed that OCI showed fastest migration. (c, d, e) Plots represent means ± sem. Statistical analyses: (c, d, e) one-way ANOVA followed by Dunnett’s test for multiple comparisons. Multiple comparisons: (c, d, e) OCI vs. MDA; OCI vs. HSPC.

We found that migration patterns, distance, and speed of migration exhibited tumor-and cell-type specific behavior. Specifically, breast tumor cells moved relatively shorter distances compared to leukemic cells or hHSPCs. Further, leukemic cells showed faster intravascular dissemination and preferred to stay in circulation (depicted by straight lines in the 2D tracking map). Solid breast tumor cells, however, tended to adhere to one site, specifically to the caudal hematopoietic tissue (CHT) of the zebrafish embryo. Mixed behavior was observed with the hHSPCs with some cells adhering to the CHT and the rest persisting in circulation (depicted by straight lines), as shown by the 2D tracking map (Fig_2b). Concurring with visual observations from SPIM movies, we found that a leukemic cell covered a significantly longer maximum distance travelled in anterior-posterior and dorsal-ventral direction (459.0 ± 70.05 μm) compared to breast tumor cells (91.44 ± 6.08 μm; P < 0.0001) or hHSPCs (83.55 ± 9.58 μm; P < 0.0001) (Fig_2c). Moreover, leukemic cells displayed a significantly higher net distance (in anterior-posterior and dorsal-ventral direction), with a mean value of 353.5 ± 58.21 μm compared to solid breast tumor cells (71±5.25 μm; P < 0.0001). As expected, net distance for hHSPCs was similar to that of the leukemic cells (328±42.63 μm, P = 0.8453, Fig_2d). In addition, the total distance travelled was higher for hHSPCs compared to leukemic, or breast tumor cells (Suppl_fig_1). Next, we calculated the intravascular speed to be 193.6 ± 44.09 μm/s for the leukemic cells, 63.6 ± 7.01 μm/s for breast tumor cells (P < 0.0001), and 83.55 ± 9.57μm/s for hHSPCs (P = 0.0008), indicating that the leukemic cells travelled significantly faster than breast tumor cells or hHSPCs (Fig_2e).

### Metastatic breast tumor cells can actively migrate and survive longer within the host

After injection, tumor cells generally migrated in the direction of blood flow as they were injected in the DoC. In order to discriminate whether tumor cells disseminated passively due to blood flow or migrated actively, we used silent heart morpholino^31^-injected zebrafish for xenotransplantation. In this no-blood-flow environment, we observed that even though most of the injected metastatic breast tumor cells remained at the site of administration, less than 10% of the cells migrated to the tail region (Suppl_fig_2), indicating that these cells are capable of active migration.

Next, to investigate whether the observed distribution is a specific feature of malignant cells, we xenotransplanted and traced malignant breast tumor cells (MDA-MB231) and their non-malignant counterparts, breast epithelial cells (MCF10A), in *casper* embryos. As reported previously^20^, much lower numbers of normal breast epithelial cells survived compared to breast tumor cells over time (Fig_3a), irrespective of whether these cells were located in the head, trunk, or tail regions (Fig_3b). Quantification of cell numbers at 4 dpi revealed that the highly metastatic breast tumor cells persisted much longer in the host (53.78 ± 1.74%, P < 0.0001, for all regions), whereas the fluorescence signal of normal epithelial cells regressed after 48 hours of transplantation (16.24 ± 0.99%), indicating engraftment failure of the non-malignant cells (Fig_3c, 3d, 3e).

**Fig_3.**
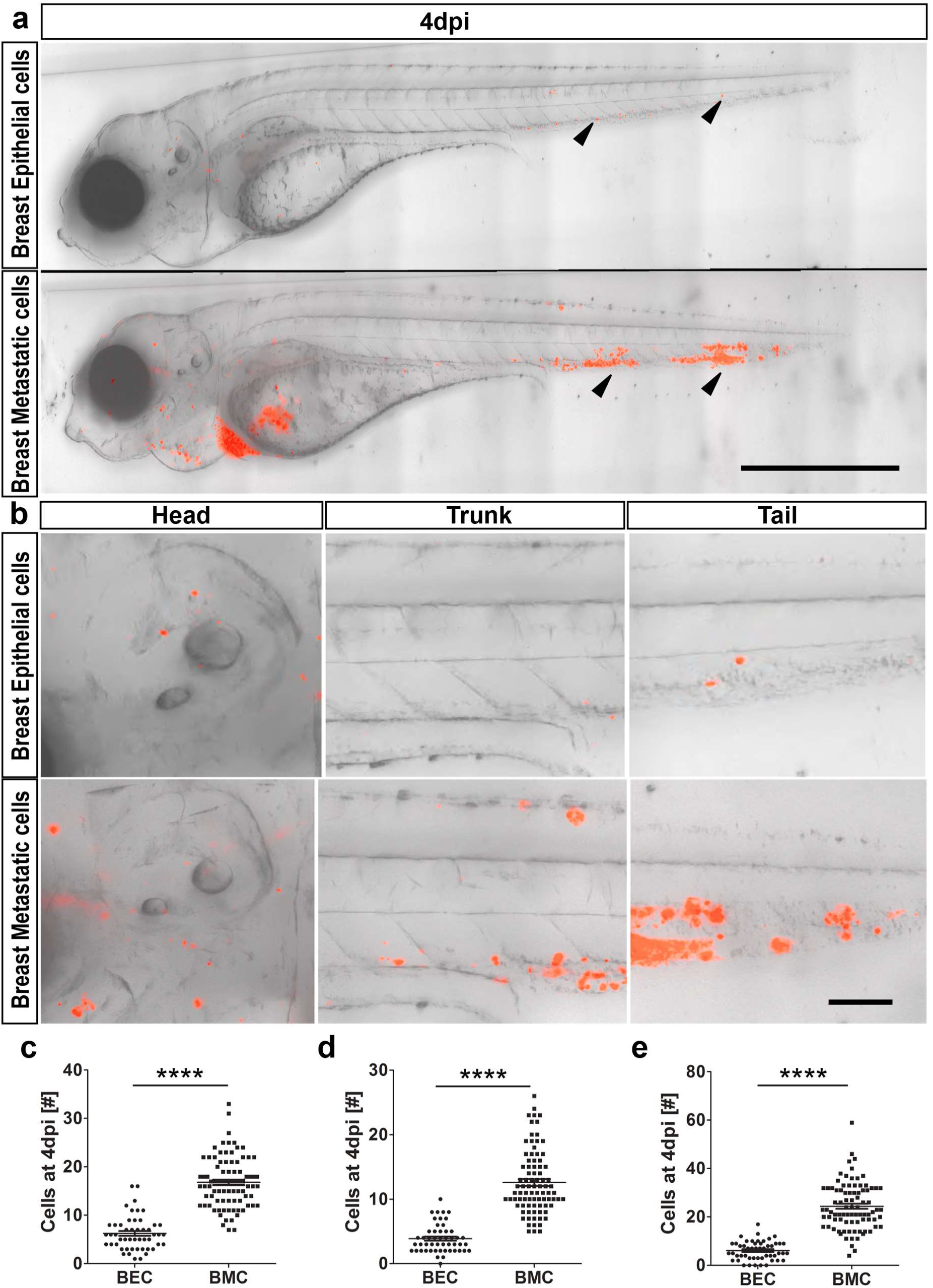
Dissemination of epithelial versus metastatic cells. (a) At 4 dpi, breast epithelial cell (BEC) numbers were drastically reduced compared to breast metastatic cells (BMC), as indicated by black arrowheads. Cells were depicted in red on a transmission image of zebrafish. Scale bar 500 μm. (b) Representative images of head, trunk, and tail regions of both BEC and BMC with high magnifications. Scale bar 50 μm. (c, d, e) Quantifications of the cells at 4 dpi in all the regions. Head (c), trunk (d), and tail (e) showed that breast metastatic cells survived better in the eZXM. In all the regions observed, the cell numbers were significantly higher for breast metastatic cells. (c, d, e) Plots represent means ± sem. Statistical analyses: two-tailed Mann-Whitney’s U-test.

### Host immune cells react to injected tumor cells

Cell numbers of injected metastatic breast tumor cells showed a tendency to decline from 1 dpi, with significant reduction to about 40-50% of the original cell number at 3 dpi, suggesting a reduced survival after xenotransplantation (Suppl_fig_3). Interestingly, high magnification SPIM time-lapse recordings revealed a host cell enclosing a tumor cell (Suppl_movie_4). Therefore, to test whether the host immune system is responsible for the observed reduction in injected tumor cells over time, we investigated the role of immune cells, specifically macrophages, by injecting metastatic MDA-MB231_eGFP (breast tumor) cells into the *Tg(mpeg1:mCherry)* zebrafish reporter line, in which macrophages are labeled by mCherry^32^. We observed co-localization of tumor cells with macrophages (Fig_4a), and quantification of the number of double positive cells (macrophage co-localizing with a tumor cell) over time showed a significant increase in co-localized cells at 3 dpi (4.37±0.80 cells) compared to 24 hpi (1.70±0.42 cells; P=0.0103), thereby confirming a potential role for innate host immune cells in restricting the survival of injected human tumor cells (Fig_4b).

**Fig_4.**
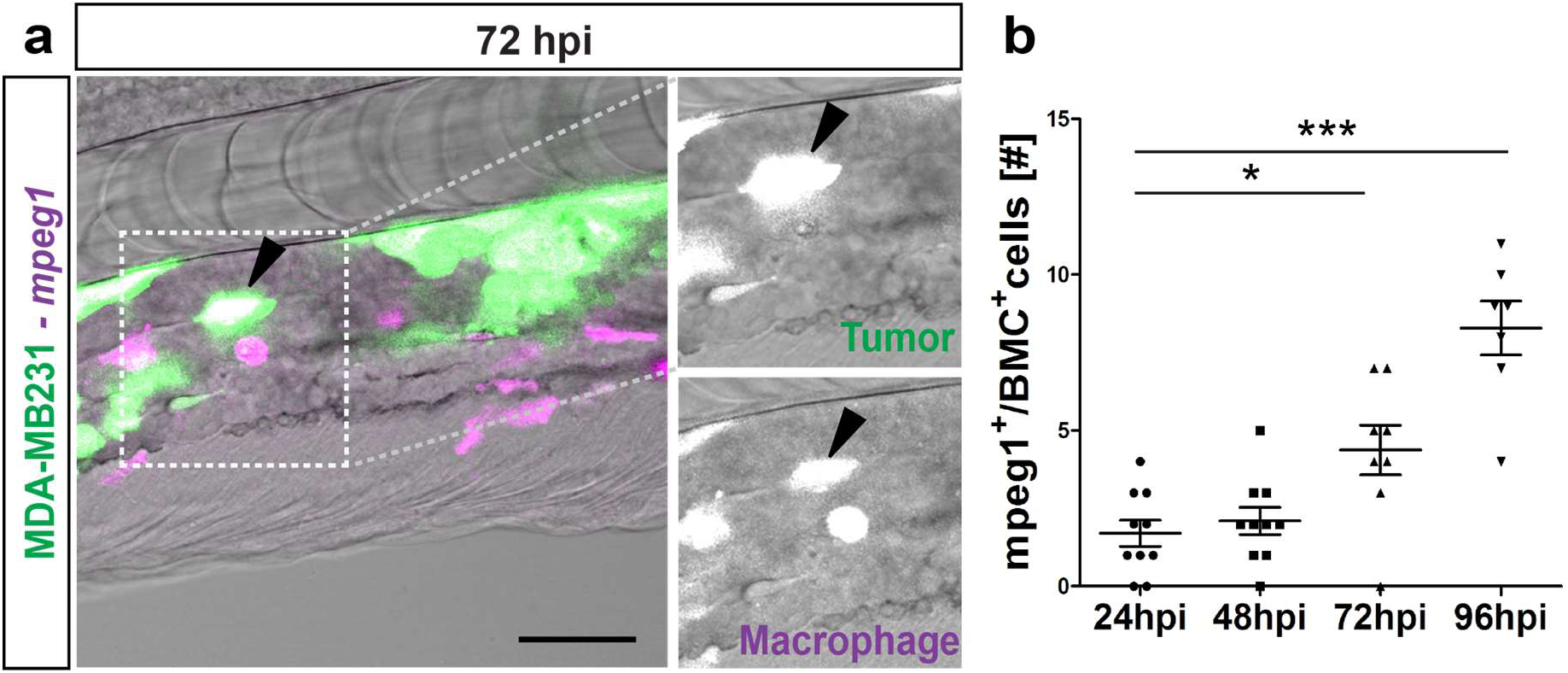
Macrophages react to tumor cells. (a) Representative image of GFP-labeled breast metastatic cells (BMC) (MDA-MB231) xenografted in eZXM expressing mCherry-labeled macrophages. Co-localization of tumor cells with macrophages was observed (black arrowhead). Higher magnification of boxed region with MDA-MB231 cell (green – top) and macrophage (magenta – bottom). Scale bar 100 μm. (b) Quantification of the co-localization of tumor cells (BMC+) with macrophages (mpeg1+) over time. At 72 hpi, a significant increase in co-localization was observed. Plot represented means ± sem. Statistical analyses: one-way ANOVA followed by Dunnett’s test for multiple comparisons. Multiple comparisons: 24 hpi vs. 48 hpi (P = 0.9225); 24 hpi vs. 72 hpi (P = 0.0103); 24 hpi vs. 96 hpi (P < 0.0001).

### Extravasation and caudal tail invasion by metastatic tumor cells

We examined whether extravasation and tissue invasion by tumor cells, the hallmarks of tumor metastasis, could be observed in real-time using the eZXM. To visualize extravasation, we acquired high magnification time-lapse images of the tail region of the eZXM after they were injected with metastatic MDA-MB231 cells at the DoC. An extravasation event was recorded when tumor cells were found leaving the vessels and entering into the surrounding tissue, and high-resolution images provided a unique insight into the process of extravasation (Suppl_movie_5). Specifically, tumor cells appeared to change shape from ‘round’ to one with extending protrusions. After a few hours, these protrusions extended their filopodia-like structures into the tissue near the vessel wall. We speculate that the cells with protrusions near the vessel wall mark the initiation of the extravasation event. Additionally, confocal images of fixed embryos at 4 dpi revealed that these protruding cells near the vessel wall pushed their entire cellular contents outside the vessel wall, marking the completion of extravasation, as shown in Fig_5a.

**Fig_5.**
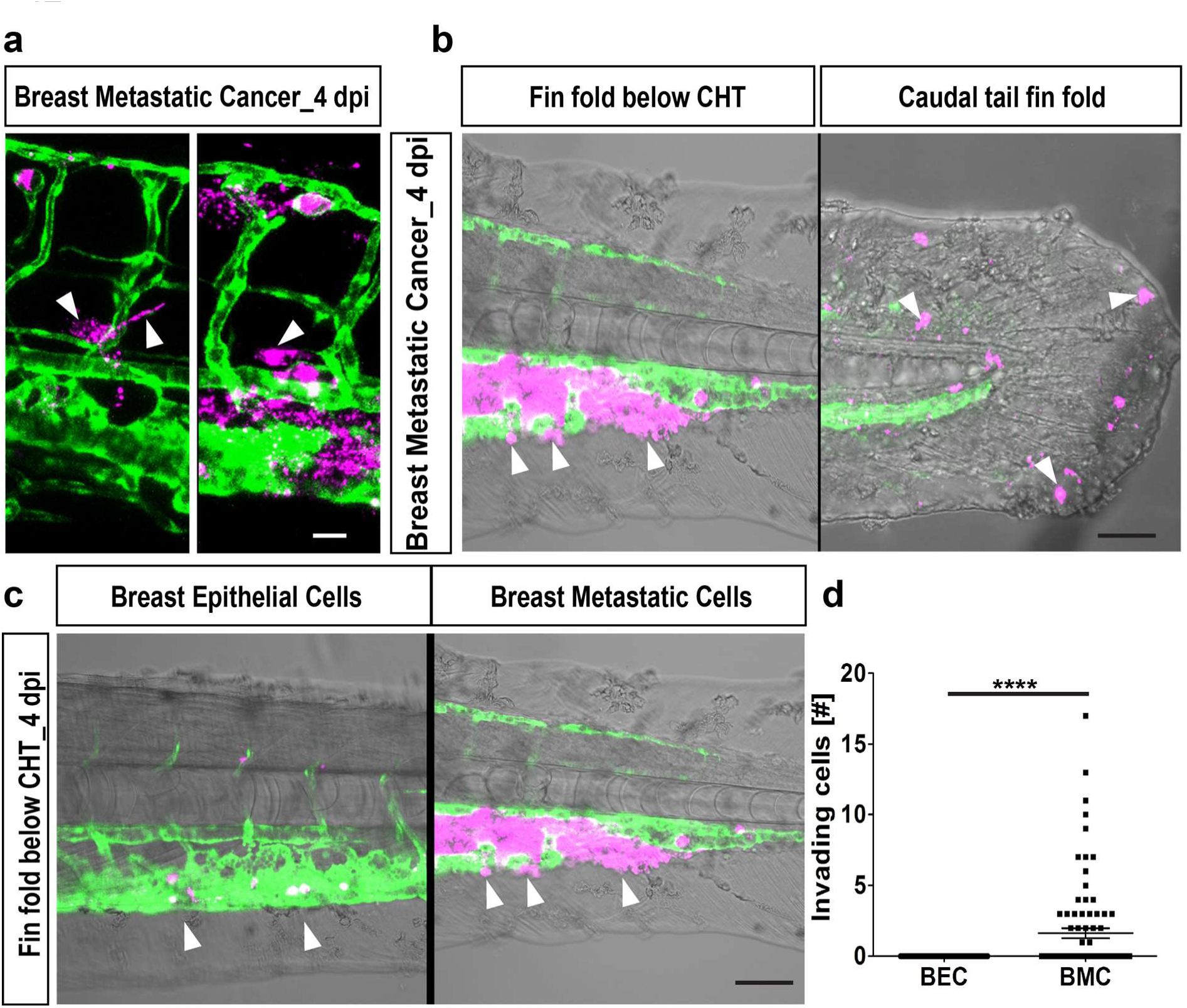
Extravasation and invasion of tumor cells. (a) Representative images of breast cancer cells initiating extravasation by forming protrusions (left, white arrowheads). These protruding cells later pushed the entire cellular content into the surrounding tissue (right, white arrowhead). Scale bar 20 μm. (b) Breast metastatic cells invaded the avascular tail region. Two representative images of invading tumor cells (white arrowheads) in the fin-fold below the CHT region (left) and caudal tail fin-fold (right). Scale bars: 150 μm. (d) Quantification at 4 dpi revealed significant tail invasion of the breast metastatic cells (BMC) over breast epithelial cells (BEC). Plots represent means ± sem. Statistical analyses: two-tailed Mann-Whitney’s *U*-test.

Next, we also assessed invasion behavior. The injected tumor cells were termed invading cells^20^ if they migrated outside the vasculature and into the avascular caudal tail region. At 4 dpi, metastatic MDA-MB231 cells appeared to invade the avascular fin-fold below the CHT and into the caudal fin fold (Fig_5b). Importantly, metastatic breast tumor cells were capable of invading the avascular tail fin while normal breast epithelial cells did not, thus displaying characteristic tumorigenic behavior (Fig_5c). Invasion at the caudal tail fin at 4 dpi was observed in 31.81% (28/88 injected embryos) of the MDA-MB231 injected embryos (Fig_5d) compared to 0% in MCF10A injected embryos (0/63 injected embryos; P < 0.0001). Further, primary tumor cells from breast cancer patients invaded the tail fin as early as 1 dpi in the eZXM (Suppl_fig_4).

### Rock-inhibition inhibits leukemic cell dissemination

To evaluate the utility of the eZXM for drug screening, we studied the effects of the ROCK1 (Rho-associated coiled-coil protein kinase 1) inhibitor Fasudil in our model using leukemic tumor cells. As our colleagues have demonstrated that ROCK1 inhibition suppresses leukemic growth in-vitro and engraftment in a long-term murine xenotransplantation model^33^, we set out to study its effects in our in *vivo* set-up. We found that, compared to the untreated control embryos at 24 hpi (82.67 ± 11.02), Fasudil-treated embryos showed a significant reduction in tumor cell numbers (42.36 ± 5.14, P = 0.0019; Fig_6a, Fig_6b). Moreover, to validate SPIM as a screening platform, we performed a pilot experiment wherein leukemic tumor cells were co-injected with 30 μΜ Fasudil in eZXM embryos. Both the control and Fasudil co-injected embryos were imaged simultaneously. Visual comparison of the control and Fasudil-treated embryos suggested that Fasudil indeed decreased leukemic tumor cell survival in zebrafish (Fig_6c). Quantification of all cells inside the xenografted embryos (cells inside the yolk were excluded) at the start of the experiment (0 h) and after 12 h showed that 72% ± 0.4% (mean ± SEM, n=2) of all cells survived in controls, while only 45% ± 4% (mean ± SEM, n=3; P = 0.025) survived in Fasudil co-injected embryos (Fig_6d). These observations imply that the eZXM model described here could be used as functional screening platform to identify compounds interfering with tumor cell spread and invasion.

**Fig_6.**
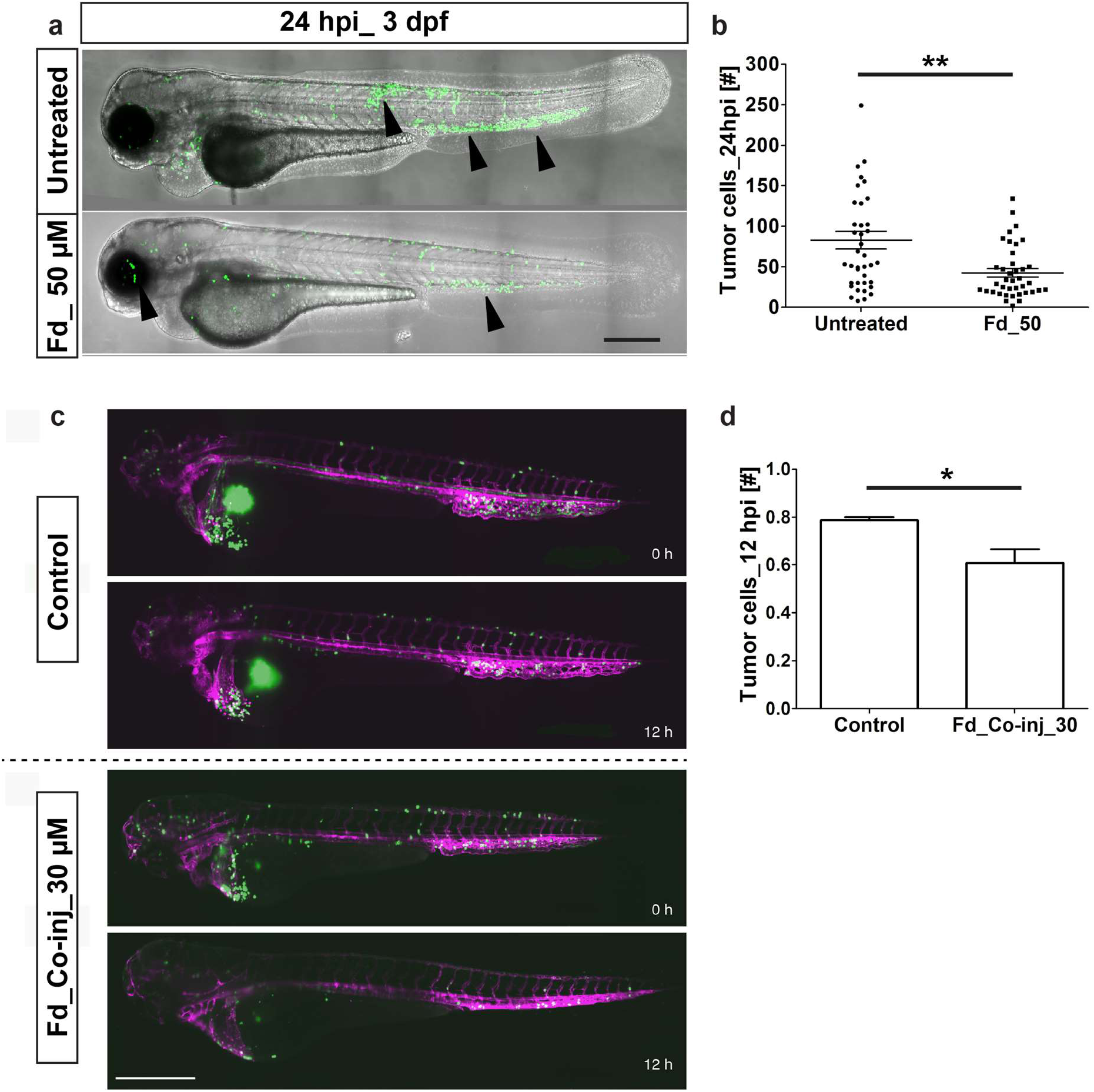
Effect of Fasudil on leukemic cells in the eZXM. Validation of the eZXM with the ROCK1 inhibitor Fasudil. (a) 50 μΜ Fasudil treatment (bottom) of the eZXM revealed a significant reduction in eGFP labeled leukemic cells (black arrowheads) at 24 hpi compared to untreated controls (top). (b) Quantification of tumor cells at 24 hpi showed a decrease in tumor cell number in 50 μΜ Fasudil-treated embryos. (c) Frames from the time-lapse movies showed dissemination of eGFP-tagged leukemic cells inside the eZXM expressing the vasculature marker *Tg(kdrl:Hsa.HRAS-mCherry)* (magenta) at the beginning of the experiment (0h) and after 12 hours (12h) in treated and untreated fish. (d) Quantified survival rate of the leukemic cells as observed from the in *vivo* time-lapse movies (right). Scale bar 500 μm. (b, d) Plots represent means ± sem. Statistical analyses: (b) two-tailed Mann-Whitney’s U-test, (d) Welch Two Sample t-test.

## Discussion

Over the past decades, different techniques have been developed to understand the mechanisms underlying the process of metastasis as it is the major determinant of cancer-related morbidity and death. The eZXM, as reported by others previously^14,15,20,23,34,35^, is a useful model to investigate metastasis in real time. Here, we describe how SPIM, a rapid, efficient, and non-invasive imaging approach^36^, can be used to visualize and understand the metastatic cascade in *vivo.* We chose SPIM as it captures the spatiotemporal dynamics of individual tumor cells in circulation with high precision and resolution, and used it to continuously track the *in vivo* behavior of two different metastatic cell lines and healthy hHSPCs in the eZXM. Despite tumor cells and hHSPCs spreading throughout the embryo due to passive dissemination via blood flow, MDA-MB231 tumor cells also migrated actively in the absence of blood flow (silent heart morpholino), and SPIM time-lapse images revealed acute differences in migration patterns among all tested cell types. The observed migration of the fluorescent solid tumor cells through circulation to distant sites in the embryo was similar to that reported previously (Tulotta et *al., 2016).* However, after 5 hpi, solid tumor cells reduced their movement and were predominantly adherent in the tail, especially in the CHT region. This finding might be explained by abundant CXCL-12 (a chemokine) expression in the CHT and its association with the homing of metastatic breast tumor cells^21^. In contrast, leukemic cells mostly remained in circulation, compliant with their origins, and their migration in the eZXM can best be described as ‘flying’ due to the swift intravascular migration; hHSPCs displayed hybrid behavior with some cells remaining in circulation and others homing to the CHT region.

High-resolution SPIM data also facilitated the analysis of tumor cell morphology. Interestingly, solid tumor cells demonstrated a change in cell shape from round to amoeboid with a protruding arm, which helped their migration from the dorsal longitudinal anastomotic vessel to the intersegmental vessels. This phenomenon also explains the nature of tumor cells to adapt to their environment to stay motile^37^. Distinctly, neither the leukemic cells nor the hHSPCs showed this amoeboid and protruding nature. Further, when a large number of solid tumor cells (~500 cells) were injected into the DoC, time-lapse images revealed huge protrusions and network like formations between neighboring cells, probably to establish stable homotypic contacts that are crucial to promote collective migration and tumor progression (unpublished data/data not shown).

Quantitative analysis of tumor dissemination in the eZXM after transplantation and related aspects including total number of tumor foci, tumor cell burden, average and cumulative distance traveled from the injection site have been previously characterized^34,38^. Many investigators have also used intravital microscopy to measure these parameters in *vivo* in mouse tissue after invasion^39^. Nevertheless, continuous in vivo monitoring of the cells and associated changes in quantitative parameters has not been reported, and a semi-automated, in-house developed tracking platform used to quantify SPIM observations revealed that net and maximum distance travelled were always higher for leukemic cells compared to solid breast tumor cells or hHSPCs, while the total distance travelled was higher for hHSPCs. This can be explained by a migration trend, which is a combination of persisting in circulation and resting in CHT sites. Also, we have calculated for the first time, the intravascular speed in anterior-posterior and dorsal-ventral direction of tumor cells in the eZXM, and as expected, leukemic cells were much faster than hHSPCs and the barely-migrating solid breast tumor cells.

Despite the immune-tolerant state of embryonic tissue, metastatic breast tumor cell numbers started to decline around 72 hpi. Tumor infiltration by macrophages during metastasis is a known phenomenon^1^, and a previous study has demonstrated that host macrophages can recognize and kill xenogenic tumor cells^40^. In concordance, high-resolution SPIM images revealed engulfment of a breast cancer cell by a perivascular host cell and increased co-localization of these tumor cells and host macrophages at 3 dpi, suggesting an indirect role for macrophages in reducing injected tumor cell numbers.

Previous reports have investigated the invasion and micrometastatic properties of xenografted cells in the caudal tail fin^23^, the dynamics of tumor cell extravasation, and associated tumor cell-endothelial cell interactions to remodel the vasculature^41^. Invasion at the caudal tail fin fold and fin folds below CHT by the metastatic breast tumor cells, but not the breast epithelial cells, attests to their tumorigenic property. Most importantly, the observed bio distribution of malignant cells could be partly confirmed using primary patient-derived tumor cells, thus supporting the relevance and potential clinical applicability of the described model. The zebrafish embryo could, therefore, complement existing patient-derived murine xenograft models.

Finally, we used Fasudil^33^, a ROCK1 inhibitor with the known effects, to validate our functional model. In line with the report of Wermke et al., 2015, we observed a 42 *%* reduction in leukemic cells in the Fasudil-treated embryos at 48 hpi. Further studies will be required to dissect the mechanistic aspects of ROCK1 inhibition during the various phases of leukemia cell spread and survival. Importantly, the intravascular distance and speed measurements reported here based on the SPIM movies can be used to provide insight into the mode of action of this drug and other such specific compounds. In contrast with the current treatment strategies involving non-specific cytotoxic drugs, we envision that our quantitative screening strategy will help screen for drugs that may interfere with adhesion and migration of metastatic tumors.

To conclude, we show that tumor cells retain their defining characteristics even after injection in to the eZXM, making the eZXM a useful screening tool. Further, tumor cell dissemination characteristics described here can be used to gain insight into mechanisms of anti-tumor action of drugs. Therefore, we propose a combinatorial approach of using the eZXM with *in vivo* SPIM imaging as a functional screening platform that can complement current drug screening and personalized anti-tumor strategies.

## Methods

### Animal care and handling

The zebrafish *(Danio rerio)* strains were kept under standard conditions (28 °C in E3 buffer) until 48 hpf as described previously^42^. All zebrafish experiments and procedures were performed as approved by the local legal authority (reference number: 24-9168.11-1/2012-32).

### Cells and cell culture

Breast tumor cells (MDA-MB231) and normal breast epithelial cells (MCF10A) were obtained from the German Collection of Microorganisms and Cell Cultures (DSMZ, Braunschweig, Germany) and from ATCC (LGC Standards GmbH, Wesel, Germany), respectively, and were cultured as described previously^43^.

We established and standardized a protocol for the isolation of primary tumor cells from breast cancer patients (n=20, tumor stages 1 and 2) using a tumor dissociation kit (130-095-929) and a tumor cell isolation kit (130-108-339) (both from Miltenyi Biotec). The isolated cells were characterized by immunofluorescence staining for pan-cytokeratin and immunophenotyping for CD24/CD44.

Isolation of CD34^+^ hematopoietic stem and progenitor cells (HSPCs) from mobilized peripheral blood (PB) obtained from healthy donors was performed using established procedures, as previously described ^44^

To obtain the human leukemic cell line OCI-AML3_eGFP, we first produced lentiviral vector particles by transfecting HEK293T cells with the lentiviral vector pRRL.SIN.cPPT.SFFV.GFP.WPRE^45^ along with the packaging plasmids psPAX and pVSVg using polyethylenimine (PEI). Lentivirus-vector containing media was collected 48 h after transfection. OCI-AML3 cells were then infected with lentiviral vector particles (0.5 x viral supernatant) in the presence of 1 mg/mL protamine, GFP expression was evaluated by flow cytometry, and positive clones were sorted using the BD FACSAriaTM II cell sorter (BD Biosciences).

### Human tumor cell preparation for transplantation and microinjection

Tumor cells were labeled with the fluorescent cell tracker CM-DiI (a lipophilic tracer, Invitrogen). A cell suspension was prepared from a 70-80% confluent monolayer as follows. Cells were trypsinized using trypsin-EDTA (0.05%), washed once in complete medium, centrifuged at 1,200 rpm for 8 min, re-suspended in PBS, and 4 μl of CM-DiI added. The cell suspension was incubated in a 37 °C water bath for 4 min, immediately transferred to ice for 20 min, centrifuged for 5 min at 1,200 rpm, and the cell pellet suspended in transplantation buffer at 100-150 cells/1 nl. *Casper* embryos (45hpf) were manually dechorionated and anesthetized using 0.02% tricaine and transferred to a petri dish containing 1.5% low melting agarose in E3. Tumor cells tagged with CM-DiI were loaded in a glass capillary and micro-injected into the blood circulation of multiple zebrafish lines *(casper^11^, [Tg(kdrl:EGFP)^s843^]^29^* and *Tg(kdrl:Hsa.HRAS-mCherry)^46^)* via the duct of Cuvier (DoC). Engrafted embryos were maintained in a new petri dish at 33 °C. Based on the fluorescence spread of the injected embryos at 2 hours post injection (hpi), embryos with tumor cells in the blood circulation were selected for experiments.

### Image acquisition and processing

In order to analyze migration of injected tumor cells, live imaging of the engrafted embryos was carried out using multidirectional selective plane light sheet illumination microscopy (mSPIM)^30^. Injected embryos were imaged with SPIM for approximately 30 h at 7x magnification with images acquired every 10 min. For high-resolution imaging of extravasation, only the tail portion of the embryo was imaged with 14x magnification. To ensure constant temperature, a perfusion chamber, maintaining 33 °C, was installed. Images were later stitched and processed using an in-house developed Image J plugin. Sample drift was corrected with a rigid registration using the image registration software elastix^47,48^. For quantification, embryos were fixed in 4% paraformaldehyde at 4 °C overnight. Fixed embryos were imaged using inverted confocal microscopy (Zeiss LSM 780) at 20x magnification (whole embryos) or at 40x magnification (tail region). Confocal stacks were converted to maximum intensity projections using Image J (v 1.51h).

### Tracking analysis

An in-house developed tracking method was used to analyze the maximum intensity projections of the time-lapse images generated by the mSPIM. This semi-automated tracking analysis combines three already existing and broadly-used open-source software tools, namely CellProfiler (v. 2.1.1) ^49^, CellTracker^50^ and R (v. 3.1.2; CRAN; R Core Team, 2014). CellProfiler was used for image segmentation as well as for an automated pre-tracking step. The resulting data was thereafter transcribed into a .xml file using R. Potential mistakes that occurred during the automated tracking process could then be corrected with the help of CellTracker, before the data was finally analyzed and visualized using R. Several measurements could be extracted from the resulting migration data. In the presented work, we concentrate on the maximum distance between any two points in the migration path, the net distance between the final position and the injection site (origin), the total distance travelled as well as the migration speed of the cells.

### Drug Treatment and Efficiency Evaluation

Functional validation of the model was carried out using the Rock-inhibitor, Fasudil^33^. A working concentration of 50 μΜ Fasudil in water, prepared from 1 mM stock, also in water, was added to E3 medium containing injected embryos. Tumor cell survival at 24 and 48 hours post injection (hpi) was assessed by fluorescence microscopy.

### Statistical analysis

Statistical analysis was performed using the Prism software (Ver.6.0, GraphPad La Jolla, USA). Results are expressed as the mean +/-SEM. If not indicated otherwise, student’s t-test or one-way analysis of variance (ANOVA) were performed followed by the Dunnett’s method for multiple comparisons. P < 0.05 was considered to be statistically significant (*0.01 < P < 0.05; **0.001 < P < 0.01; ***0.0001 < P < 0.001; ****P < 0.0001).

## Acknowledgements

This work was supported by the Deutsche Forschungsgemeinschaft DFG (Center of Excellence, Center for Regenerative Therapies, www.crt-dresden.de and the Collaborative Research Grant SFB 655 to MB). In addition, the authors express thanks to the light microscope facility of the CRTD, thanks to the fish facility (Biotech & CRTD) for excellent support, and Dr. Vasuprada Iyengar for editing. Furthermore, this work was supported by the German Consortium for Translational Cancer Research (www.dktk.dkfz.de) and the National Center for Tumor Diseases (www.nct-dresden.de), Dresden.

## Supplementary Figures

**Suppl_fig_1.**
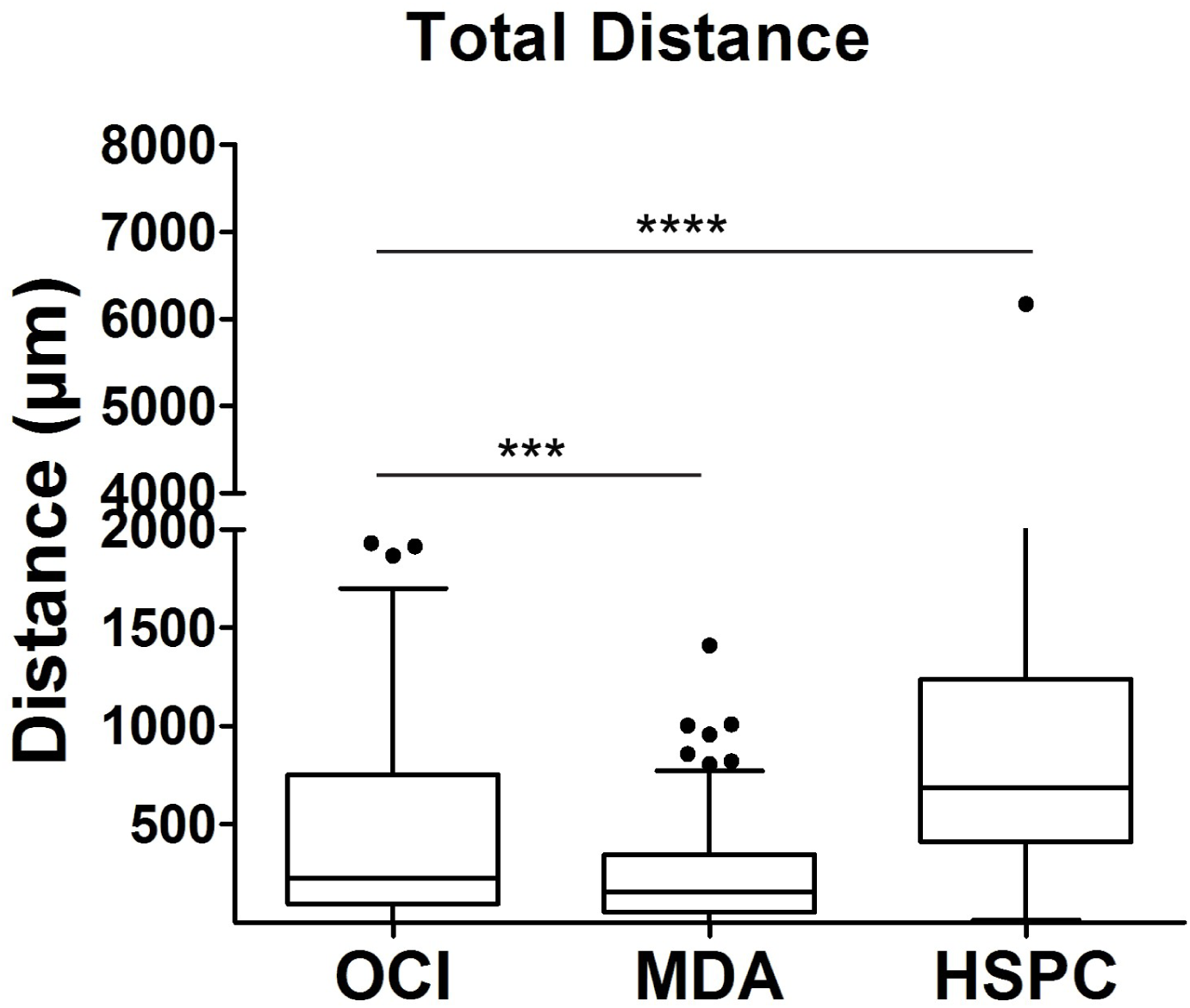
hHSPCs traverse greater total distance. The tracking measurements revealed the total distance covered by cells inside the eZXM. hHSPCs covered a significantly greater total distance compared to breast tumor cells (MDA) and leukemic cells. Plot represented means ± sem. One-way ANOVA followed by Dunnett’s test for multiple comparisons. Multiple comparisons: OCI vs. MDA (P = 0.0002); OCI vs. HSPC (P < 0.0001).

**Suppl_fig_2.**
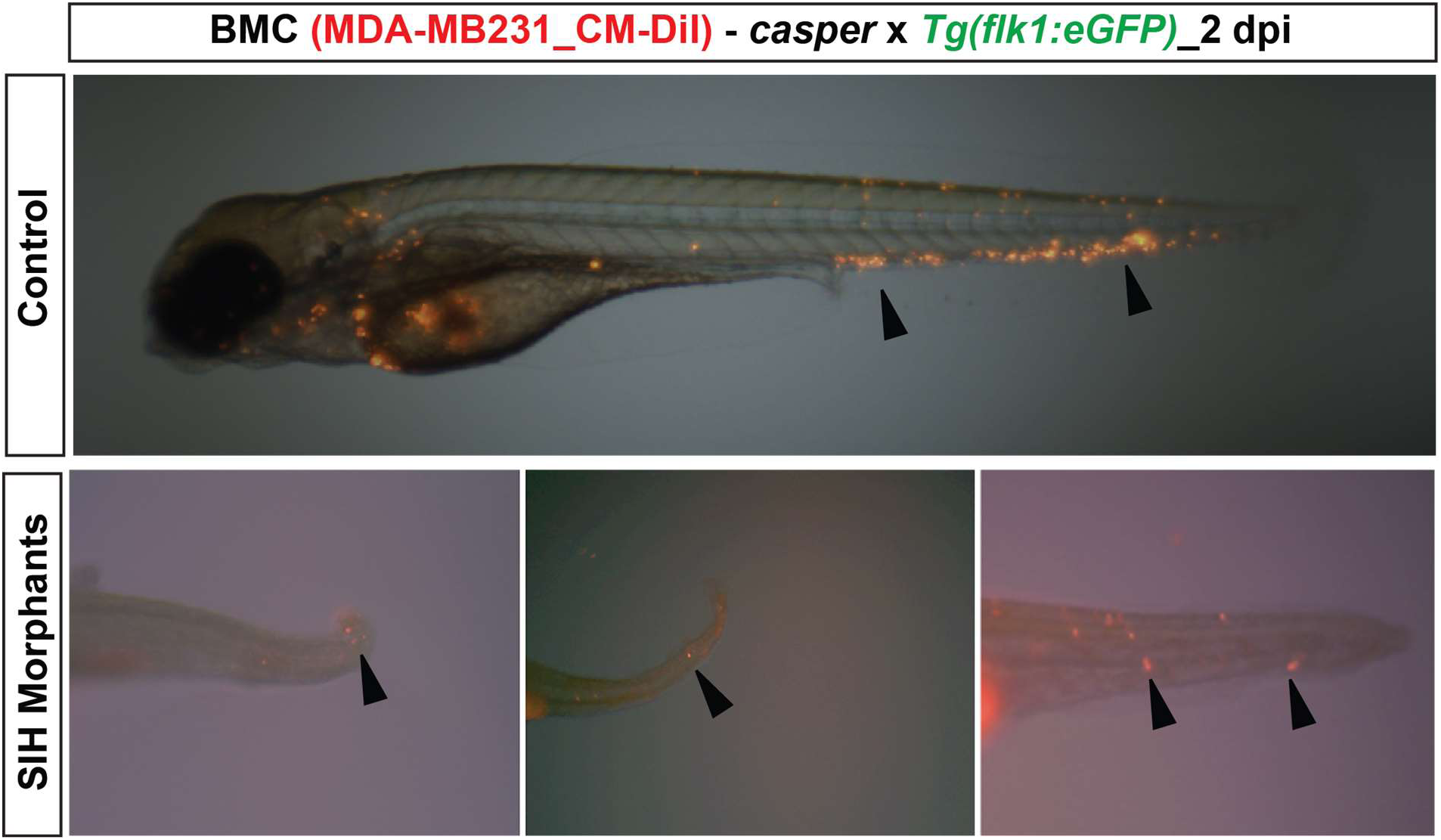
Active migration of the tumor cells. Breast metastatic tumor cells (BMC) labeled with CM-DiI were injected into silent heart morpholino injected eZXM. Silent heart morpholino stopped the heartbeat. Representative image of control morpholino eZXM injected with breast tumor cells (black arrowheads) with dissemination throughout the embryo (top). In the no-flow environment, breast tumor cells (black arrowheads) migrated actively from the site of administration towards the tail region (bottom).

**Suppl_fig_3.**
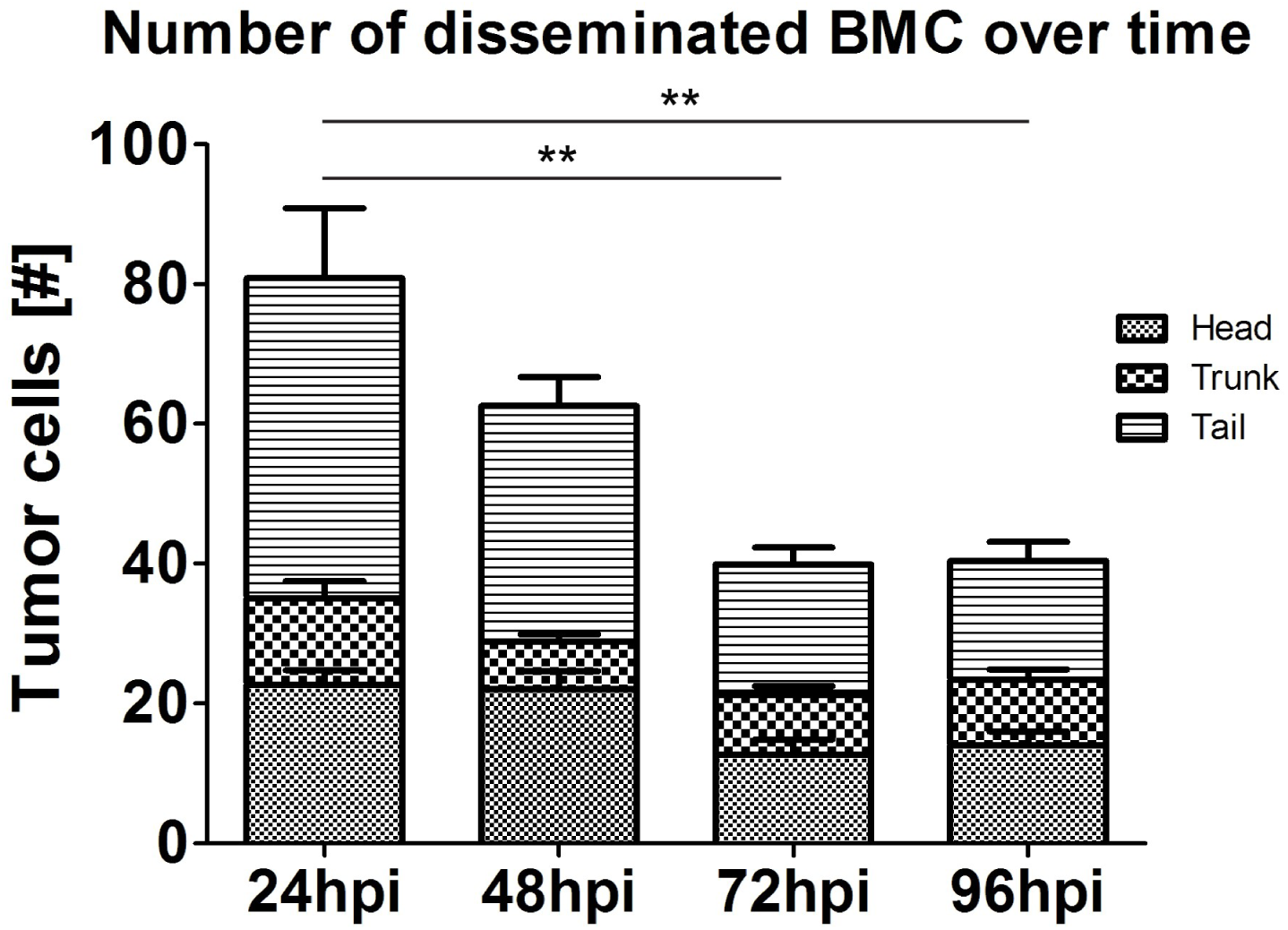
Dissemination of metastatic tumor cells over time. Breast metastatic tumor cells (BMC-MDA-MB231) were injected into the eZXM and the number of surviving cells were quantified over time. Quantification revealed that at 72 hpi, breast tumor cell numbers were significantly reduced compared to at 24 hpi. Plot represented means ± sem. Statistical analyses: one-way ANOVA followed by Dunnett’s test for multiple comparisons. Multiple comparisons: 24hpi vs. 48hpi (P > 0.9999); 24hpi vs. 72hpi (P = 0.0028); 24hpi vs. 96hpi (P = 0.0086).

**Suppl_fig_4.**
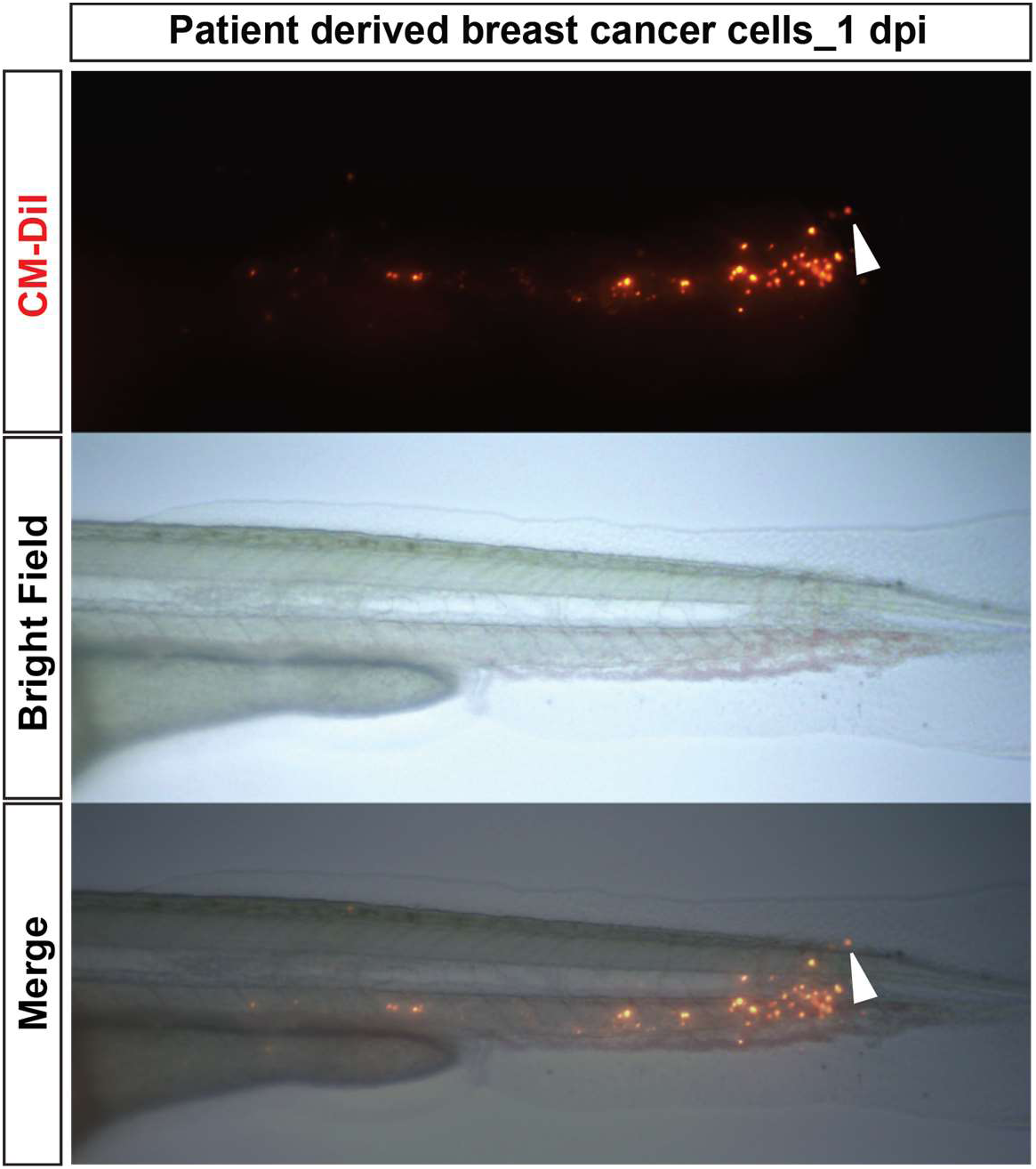
Tail invasion in the eZXM injected with primary tumor cells. Primary tumor cells (top) were labeled with CM-DiI and injected in the eZXM. Imaging of the tail part of at 1 dpi (bright field image, middle), revealed invasion of the tail fin fold region by a single cell (white arrowhead).

## Supplementary movies

### Suppl_movie_1 Dissemination of leukemic cells

OCI-AML3_eGFP cells (leukemic cells; magenta) were injected into the eZXM expressing the vasculature marker *Tg(kdrl:Hsa.HRAS-mCherry)* (depicted with green). Leukemic cells were migrating back and forth and mostly preferred to stay in circulation. Time shown as h:min:sec. Scale bar: 500 μm.

### Suppl_movie_2 Dissemination of breast metastatic tumor cells

MDA-MB231 (breast tumor cells) labeled with CM-DiI (magenta) were injected into eZXM expressing the vasculature (green) marker *Tg(kdrl:EGFP*)^s843^. After injection, the tumor cells migrated along with the blood flow, disseminated from head to tail, migrated towards the circulatory loop end, and adhered near the caudal hematopoietic tissue (CHT) region. Time shown as h:min:sec. Scale bar: 500 μm.

### Suppl_movie_3 Dissemination of human hematopoietic stem cells (hHSPC)

HSPCs labeled with CM-DiI (magenta) were injected into eZXM expressing the vasculature (green) marker *Tg(kdrl:EGFP*)^s843^. HSPCs migrated similarly to breast tumor cells and leukemic cells and were disseminated throughout the embryo. Some cells stayed in circulation while the rest of them adhered near the CHT, tail region. Time shown as h:min:sec. Scale bar: 500 цш.

### Suppl_movie_4 Host-cell enclosing a tumor cell

Metastatic breast tumor cells (green) were injected in eZXM expressing the vasculature marker *Tg(kdrl:Hsa.HRAS-mCherry)* (magenta). A tumor cell interacted with a host cell (magenta, white arrowhead). The host cell enclosed the tumor cell at 6 h after establishing contact. Time shown as h:min:sec. Scale bar: 300 μm.

### Suppl_movie_5 3D rendering of extravasation initiation

Breast metastatic cells (MDA-MB231_eGFP, in green) were injected in eZXM expressing the vasculature marker *Tg(kdrl:Hsa.HRAS-mCherry)* (magenta). 3D rendering of time-lapse SPIM movie showed a cell having a tendency to extravasate by forming protrusions (white arrowhead). Protrusion projected their arms into the surrounding lumen marking the initiation of extravasation event.

